# EPS8 dampens the growth dynamics and prolongs the lifetime of actin-based protrusions

**DOI:** 10.64898/2026.04.27.721156

**Authors:** Alexandra G. Mulligan, Zachary J. Lehmann, Kianna L. Robinson, Matthew J. Tyska

## Abstract

Actin-based membrane protrusions such as filopodia, microvilli, and stereocilia support a range of cell functions, from nutrient absorption to mechanosensation. In each case, membrane deformation is supported by a core bundle of actin filaments, organized in a unipolar ‘barbed-end out’ manner. Although their structures and proteomes are well characterized, mechanisms governing the growth and stability of these protrusions remain less clear. Factors that localize to the distal tips of these structures are of particular interest, as they are well positioned to control actin assembly at filament barbed ends. One such factor, EPS8, localizes to distal tip puncta in multiple protrusion types. While early biochemical studies suggested a role in filament capping, loss of EPS8 in multiple models shortened microvilli and stereocilia, suggesting roles in elongation. More recent studies in differentiating epithelial cells suggested that EPS8 promotes protrusion growth and stability. To clarify EPS8’s function in the distal tip compartment, we leveraged acute loss-of-function experiments and titrated gain-of-function approaches in combination with live imaging. Acute sequestration of EPS8 led to rapid depletion of filopodia. Conversely, increasing cellular EPS8 levels elevated EPS8 per distal tip punctum, increased F-actin content within individual filopodia, reduced filopodia elongation rates, increased protrusion lifetimes, and protected filopodia against cytochalasin D-induced collapse. These findings suggest that EPS8 binds filament barbed ends as a ‘leaky capper,’ slowing monomer addition while stabilizing bundles and preventing collapse. These activities are likely critical for building and maintaining the large arrays of protrusions that are assembled by diverse epithelial cell types.

## INTRODUCTION

Actin-based membrane protrusions and their core components can be traced back to the holozoan lineage and may play a role in the rise of obligate multicellularity (1,2). Putative filopodia have been identified in ancient eukaryotes, with microvilli later emerging in Urmetazoans and animal cells (2). Today, such structures are found across diverse organisms, tissues, and cell types. Well-characterized examples include filopodia, microvilli, and stereocilia, which function in adhesion and taxis, solute transport, and hearing and balance, respectively (3). Large actin-bundle supported protrusions are also found in the sperm acrosomal process and angiosperm pollen tubes, both of which facilitate fertilization (4,5). Despite their functional diversity, previous studies have identified molecular activities that are common to all protrusions, which are critical for the assembly and stability of these features. For example, actin filament bundling proteins are required to assemble a core bundle with sufficient flexural rigidity to deform the plasma membrane (6-9). Additionally, proteins that link the overlying membrane to the underlying actin core bundle provide additional stability and prevent neighboring protrusions from fusing (10,11). Finally, distal tip enriched proteins and an electron-dense “tip complex” have been proposed to physically couple the barbed ends of core bundle filaments to the highly curved plasma membrane at these sites (12-14). Proteins that localize to the tips are also well positioned to control actin dynamics underpinning protrusion growth, although the detailed functions and mechanisms of many tip targeted molecules remain unresolved.

One factor that exhibits robust, punctate localization to the distal tips of protrusions is epidermal growth factor receptor kinase substrate 8 (EPS8) (15-18). Initially identified as a phosphorylation target of EGFR, EPS8 has since been implicated in the signaling transduction that drives a wide range of physiological and pathophysiological processes (19,20). EPS8 consists of multiple structural motifs, including: (i) an N-terminal phosphotyrosine binding (PTB) domain that functions as a scaffold for protein-protein interactions in both a phosphorylation-dependent and -independent manner; (ii) an extended polyproline motif, known to interact with the SH3 domain of insulin receptor tyrosine kinase substrate (IRTKS)(17); (iii) a central SH3 domain that preferentially binds the PXXDY motif of proteins involved in GTPase and receptor tyrosine kinase (RTK) mediated signaling (21-25); and (iv) a C-terminal effector region that contains F-actin-binding domains and binds homologous inverse bin-amphiphysin-rvs (I-BAR) domain proteins, including insulin receptor substrate p53 (IRSp53/BAIAP2) (26,27).

EPS8 function in cells has been explored using multiple model systems, and several common phenotypic features have emerged. In *C. elegans*, EPS8 KO disrupts intestinal epithelial morphogenesis, leading to brush borders characterized by abnormally short microvilli, with a loss of density and rigidity (15). EPS8 KO in mouse models also leads to abnormally short and disorganized microvilli on the apical surface of enterocytes, and deformed stereocilia on the surface of mechano-sensory hair cells (28,29). In HeLa cells and fibroblasts, KD of EPS8 results in fewer filopodia, although one study in hippocampal neurons showed that loss of EPS8 increased the number of axonal filopodia (18,27). Whereas most of these findings indicate that EPS8 promotes protrusion elongation, they also point to the possibility of context dependent roles (30).

Live imaging studies have also proven informative for revealing how EPS8 impacts specific protrusion behaviors. At the apical surface of polarizing epithelial cell culture models, EPS8 puncta arrive minutes before the *de novo* growth of a microvillus (31). Following growth of the protrusion, EPS8 in conjunction with binding partner IRTKS, promotes the elongation and directional persistence of microvilli, as they glide across the apical surface early in differentiation (32). Those studies also revealed that loss of the distal tip EPS8 punctum precedes microvillar collapse, implicating EPS8 in protrusion survival (31). Collectively, these studies suggest that EPS8 plays important roles in protrusion growth, elongation, and survival on the cell surface.

Early *in vitro* work demonstrated that C-terminal fragments of EPS8 exhibit filament capping and bundling activities (26,33); the amphipathic H1 helix (a.a. 648-700 in human) was shown to cap actin filaments, whereas the H2-H5 helical lobe (a.a. 701-821 in human) exhibited filament bundling potential (33). Notably, full length EPS8 does not bind actin *in vitro*, suggesting that it dwells in an autoinhibited state, and binding partners such as Abi1 and IRSp53 may regulate its activity *in vivo* (25,27,34). However, capping and bundling activities seem to be at odds with what we know about EPS8 localization and function. For example, while the protrusion distal tip is the expected localization site for a capping protein, results from the loss-of-function studies outlined above indicate that EPS8 promotes filament elongation, rather than inhibiting polymerization and promoting filament turnover as expected for a capper (35). In addition, most well-studied actin filament bundling proteins distribute along the length of the bundles they create, rather than at the ends (6,36). This might imply that distal tip targeted EPS8 is poorly positioned to bundle filaments, although recent studies did reveal that some bundlers occupy discrete “neighborhoods” along the bundle axis. In the context of intestinal brush border microvilli, mitotic spindle positioning (MISP) selectively localizes to the rootlet segment (37). Regionalized distributions have also been reported for other actin bundlers, such as espin (ESPN), fimbrin (PLS1), advillin (AVIL), and LIM domain and actin-binding protein 1 (LIMA1) in intestinal tuft cells (38), and taperin (TPRN) in stereocilia (39). These latter findings raise the possibility that EPS8 might function as a spatially restricted filament bundler, cross-linking the barbed ends of filaments specifically in the distal tip compartment. Nevertheless, significant questions on the nature of EPS8 function in the distal tip compartment remain unanswered.

To more directly investigate the function of EPS8 in living cells, we leveraged acute loss-of-function experiments and a titrated gain-of-function approach that employed a new knock-in HeLa cell line where endogenous *EPS8* was tagged with mStayGold.

Acute sequestration of EPS8 led to a striking loss of filopodia, whereas increasing the whole cell concentration of EPS8 increased the number and F-actin content of individual filopodia, dampened filopodial dynamics, and increased protrusion lifetime. We also observed that Cytochalasin D, which caps barbed ends, displaced endogenous EPS8 from filopodial tips, whereas cells expressing higher levels of EPS8 were protected from this effect. Based on our findings, we propose that EPS8 interacts with filament barbed ends where it functions as a ‘leaky capper’, slowing the incorporation of new actin monomers, and offering long-term protection against actin bundle disassembly. These activities are likely important for stabilizing protrusions long-term on the cell surface, in support of diverse physiological functions.

## RESULTS

### Acute sequestration of EPS8 leads to a loss of filopodia

Constitutive loss-of-function (KD and KO) experiments have offered valuable clues on the role of EPS8 in actin-based protrusions (15,27-29). However, phenotypes in those systems could be limited by the compensatory action of other EPS8 family members, including EPS8L1, EPS8L2, and EPS8L3 (40). To acutely assess the impact of EPS8 depletion from the tips of filopodia, we employed a synthetic inducible dimerization system that takes advantage of the FRB and FKBP motifs, and their high affinity for the rapalog (rapamycin analog) compound (Fig. 1A) (41). We first generated FKBP- and FRB-tagged variants of TOMM20-mCherry and mStayGold-EPS8, respectively. Our expectation was that, in co-transfected cells, exposure to rapalog would lead to the rapid recruitment of mStayGold-EPS8-FRB to the mitochondrial surface (decorated with TOMM20-mCherry-FKBP), with a corresponding depletion from filopodial tips (Fig. 1A). Constructs were co-expressed in HeLa cells and spinning disk confocal (SDC) microscopy was used to perform live imaging, which enabled us to monitor the impact of rapalog addition. To internally control for differences in mosaic expression of the two constructs, we imaged and performed measurements on the same cells before and after 1 hour of rapalog treatment. Notably, rapid recruitment of EPS8 to the mitochondrial surface resulted in a striking loss of EPS8 puncta from filopodial tips (Fig. 1B), and a parallel significant reduction in the number of protrusions after 1 hour (Fig. 1C). These data clearly indicate that EPS8 distal tip puncta play a role in stabilizing filopodia on the cell surface.

**Figure 1.**
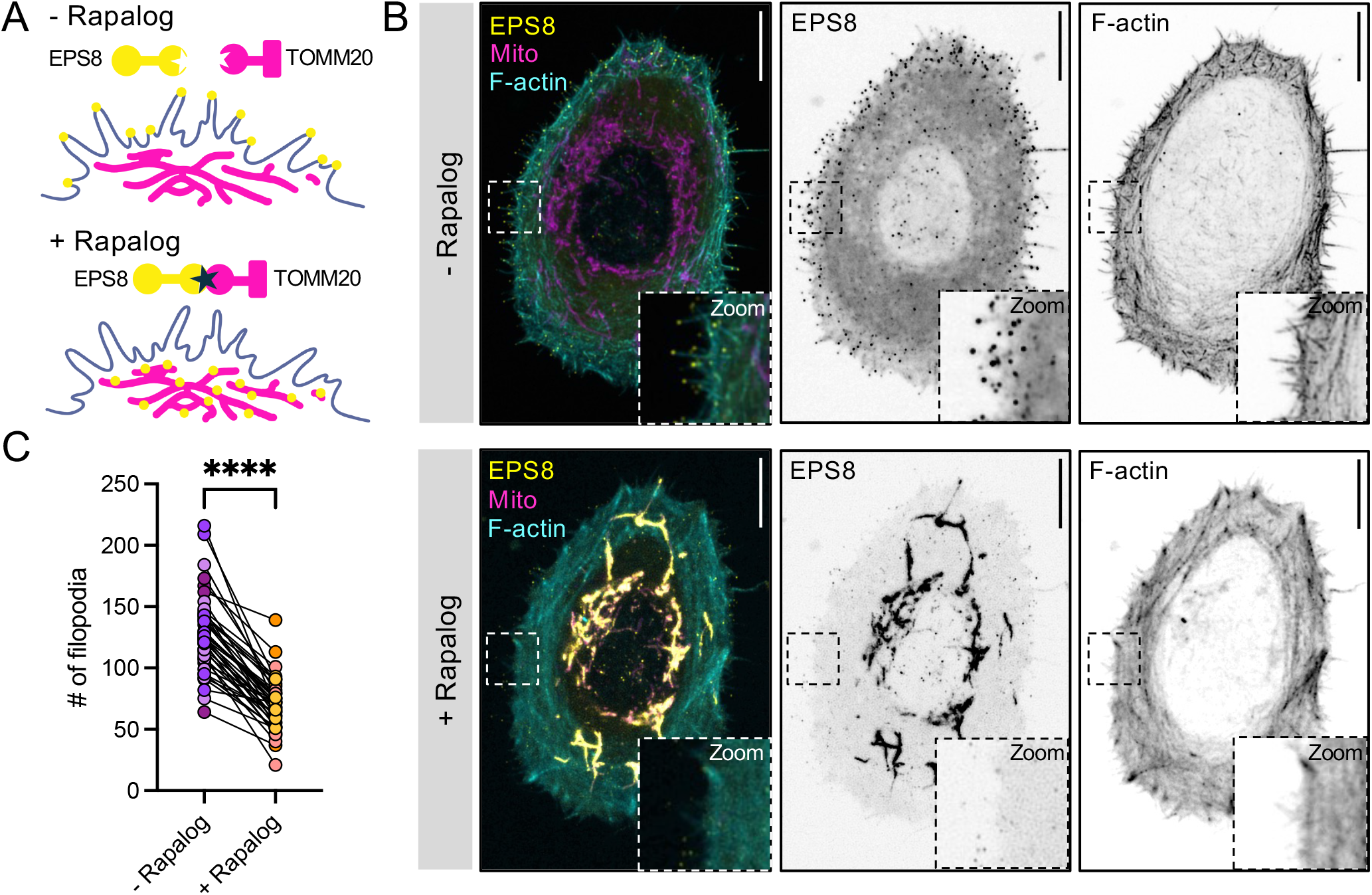
EPS8 sequestration results in loss of filopodia. **A)** Schematic depicting the rapalog-inducible system used for sequestering EPS8 to the mitochondrial surface. **B)** Max-intensity projection (MIP) 100x SDC deconvolved images of HeLa cells expressing mStayGold-EPS8-FRB-Myc (yellow) and TOMM20-mCherry-FKBP (magenta) with F-actin labeled with FastActX before and after 60 min of incubation with rapalog. Scale bar = 10 µm. **C)** Quantification of total number of filopodia post-vs. pre-rapalog induction. 45 cells measured from 3 biological replicates. Paired t-test, ****p < 0.0001.

### Raising EPS8 levels increases filopodial number and F-actin content

To complement the acute loss-of-function experiments highlighted above, we set out to examine how increasing the cytoplasmic pool of EPS8 impacts filopodial structure and dynamics. To this end, we leveraged CRISPR-Cas9 genome engineering to tag an endogenous copy of EPS8 in HeLa cells with mStayGold (Endo mSG-EPS8). As expected, filopodia on the surface of Endo mSG-EPS8 cells were uniformly decorated with distal tip puncta (Fig. 2A). To increase cytoplasmic EPS8 concentration, Endo mSG-EPS8 HeLa cells were supplemented by transient transfection with a plasmid encoding mSG-EPS8; this approach enabled us to generate and visualize cells with a broad range of EPS8 concentrations, extending up to high level overexpression (Endo + OEx mSG-EPS8; Fig. 2B,C). We focused on correlating levels of mSG-EPS8 with observed impacts on filopodia structure, F-actin content, and dynamics, from transfected vs. untransfected Endo mSG-EPS8 cells, cultured simultaneously and imaged in the same 4-chamber dish. We first examined the impact of increasing the cytoplasmic EPS8 concentration on distal tip puncta intensity. Mean puncta intensities at filopodial tips were significantly higher in cells with supplemental EPS8 vs. controls expressing only the endogenous pool (Fig. 2A,B,D). Given that EPS8 is a *bona fide* F-actin binding protein, with a proposed stoichiometry of one EPS8 molecule per filament (33), we next examined the impact of increasing EPS8 levels on F-actin content in individual protrusions. To do so, we used a live cell probe, FastActX, to label F-actin in Endo mSG-EPS8 cells with and without EPS8 supplementation (Fig. 2E). To calculate the FastActX signal density per protrusion, we generated orthogonal line scans and performed area under the curve (AUC) analysis (Fig. 2F), as described previously (42). Notably, AUC was significantly increased by EPS8 supplementation, suggesting that EPS8 positively regulates the number of filaments per protrusion (Fig. 2G). Interestingly, increasing EPS8 appeared to reduce the cortical F-actin signal associated with stress fibers, suggesting that higher levels of EPS8 promote F-actin reallocation from these contractile arrays into filopodia (Fig. S2). In addition to the F-actin signal density enrichment within filopodia, the number of protrusions per cell also increased in Endo + OEx mSG-EPS8 cells (Fig. 2H), revealing an inverse effect to depleting EPS8 from filopodial tips (Fig. 1). Together, these results indicate that raising the cytoplasmic EPS8 concentration leads to a greater number of filopodia, an increase in the amount of EPS8 per tip punctum, and more actin filaments per filopodial core bundle.

**Figure 2.**
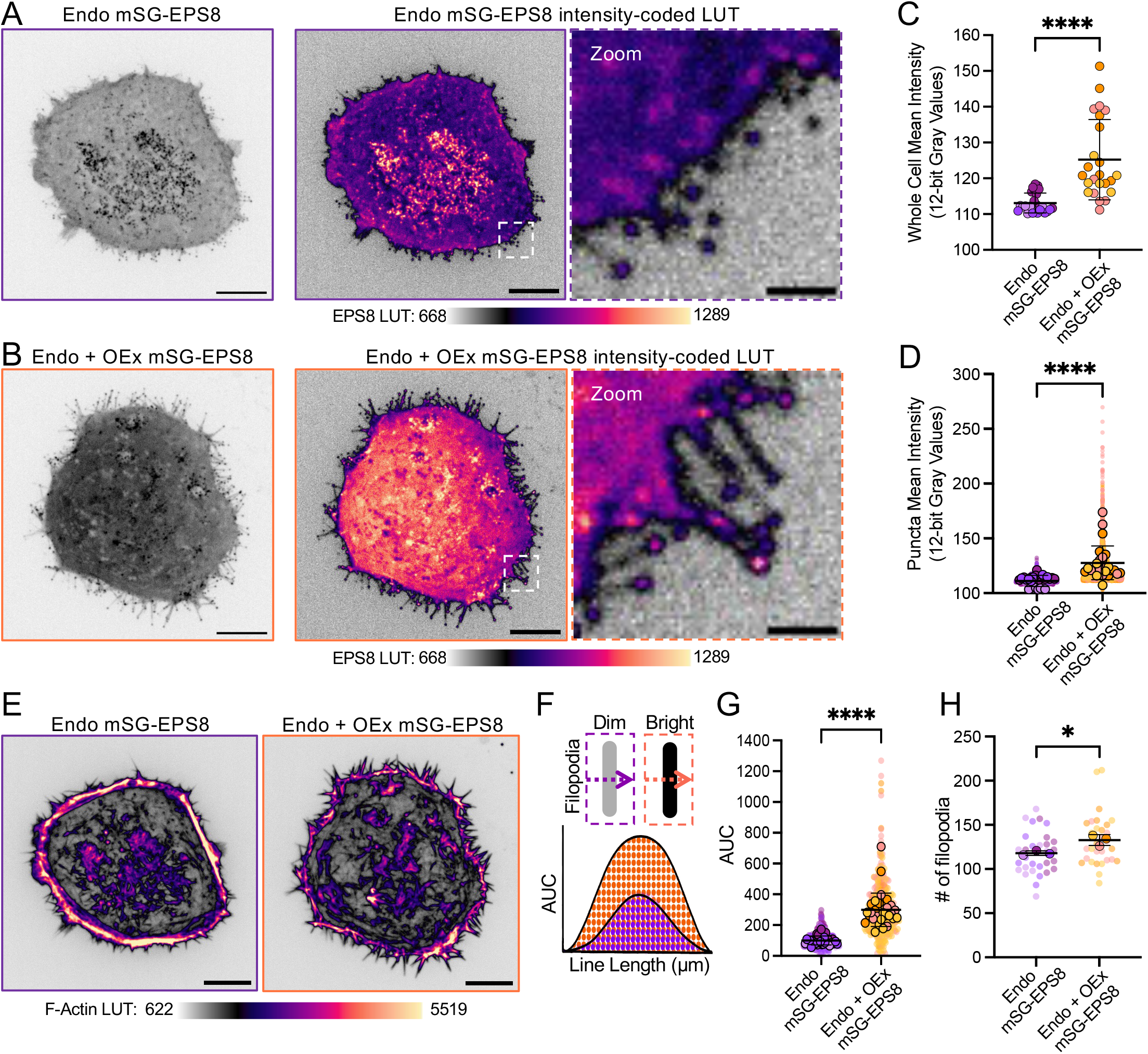
EPS8 increases protrusion number and F-actin per protrusion. **A-B)** (Left) MIP 100x SDC image of Endo-mSG-EPS8 (purple outline) and Endo + OEx mSG-EPS8 (orange outline) HeLa cells. Scale bar = 10 µm. (Middle) Intensity-matched LUT for both A & B with LUT scale. (Right) Scale bar = 2 µm. **C)** Quantification of whole cell mean 12-bit intensity values measured for Endo-mSG-EPS8 vs. Endo + OEx mSG-EPS8 cells. 25 cells measured from 3 biological replicates. Unpaired t-test, ****p < 0.0001. **D)** Superplots of mean 12-bit intensity values from EPS8 tip puncta measured for Endo-mSG-EPS8 vs. Endo + OEx mSG-EPS8 cells. Small, transparent data points show individual puncta and larger, opaque points show per cell averages. 3,405-3,457 puncta quantified over 25 cells from 3 biological replicates. Unpaired t-test on per cell averages, ****p < 0.0001. **E)** Sum-intensity projection (SIP) 100x SDC image of Endo-mSG-EPS8 (purple outline) and Endo + OEx mSG-EPS8 (orange outline) showing F-actin labeled with FastActX using intensity-matched LUTs. **F)** Schematic depicting area under the curve (AUC) analysis. **G)** Superplots of AUC measurements from filopodial FastActX signal in Endo-mSG-EPS8 vs. Endo + OEx mSG-EPS8 cells. Small, transparent data points show individual filopodia and larger, opaque points show per cell averages. 298-299 filopodia measured over 30 cells from 3 biological replicates. Unpaired Mann-Whitney t-test on the 3 biological replicates, ****p < 0.0001. **H)** Superplots showing the total number of filopodia in Endo-mSG-EPS8 vs. Endo + OEx mSG-EPS8 cells. Small, transparent data points show individual cells and larger, opaque points show replicate averages. 30 cells measured from 3 biological replicates. Unpaired t-test on the 3 biological replicates, *p = 0.0186.

### EPS8 suppresses filopodial dynamics while prolonging protrusion survival

Given that EPS8 supplementation promoted the formation of filopodia with increased F-actin content, we next sought to determine how this structural change impacted the dynamic growth properties of these protrusions. Time projected SDC movies of Endo mSG-EPS8 HeLa cultures revealed that these cells demonstrated highly dynamic edges, with cell profiles that rapidly fluctuated over time (Fig. 3A). Conversely, supplemental EPS8 appeared to attenuate these dynamics (white signal in time projection represents static or slow-moving structures; Fig. 3B). Indeed, calculating the standard deviation of cell perimeter over time revealed that cells expressing endogenous levels of EPS8 produced significantly higher values vs. cells with EPS8 supplementation (Fig. 3G), confirming that the cell periphery is more dynamic with lower levels of EPS8. To further characterize the dynamics of individual filopodia, we tracked EPS8 distal tip puncta and performed motion analysis (Fig. 3C,D). Puncta velocities in cells with EPS8 supplementation were significantly slower compared to controls with endogenous levels (Fig. 3H). Radial plots of puncta trajectories revealed that, while overall dynamics were dampened, protrusions in cells with supplemental EPS8 traveled further (Fig. 3E,F). Strikingly, the speeds of individual puncta demonstrated a hyperbolic inverse correlation to their measured EPS8 intensities, further indicating that higher levels of EPS8 directly dampen dynamics at the scale of individual filopodia (Fig. 3I). To determine if the nature of puncta movement was also impacted by EPS8 supplementation, we performed mean-squared displacement (MSD) analysis of trajectories generated from distal tip puncta tracking (43). In control cells, MSD analysis revealed that puncta at the tips of filopodia were highly mobile (Fig. 3J). In contrast, EPS8 supplementation significantly reduced mobility (Fig. 3J). Moreover, in cells with supplemental EPS8, analysis of puncta lifetimes revealed that protrusions survived approximately 60% longer vs. filopodia extending from control cells (Fig. 3K), which may contribute to the longer trajectories observed in these cases (Fig. 3F). Together, these data indicate that EPS8 dampens filopodial dynamics, while increasing the elongation and lifetime of these protrusions on the cell surface.

**Figure 3.**
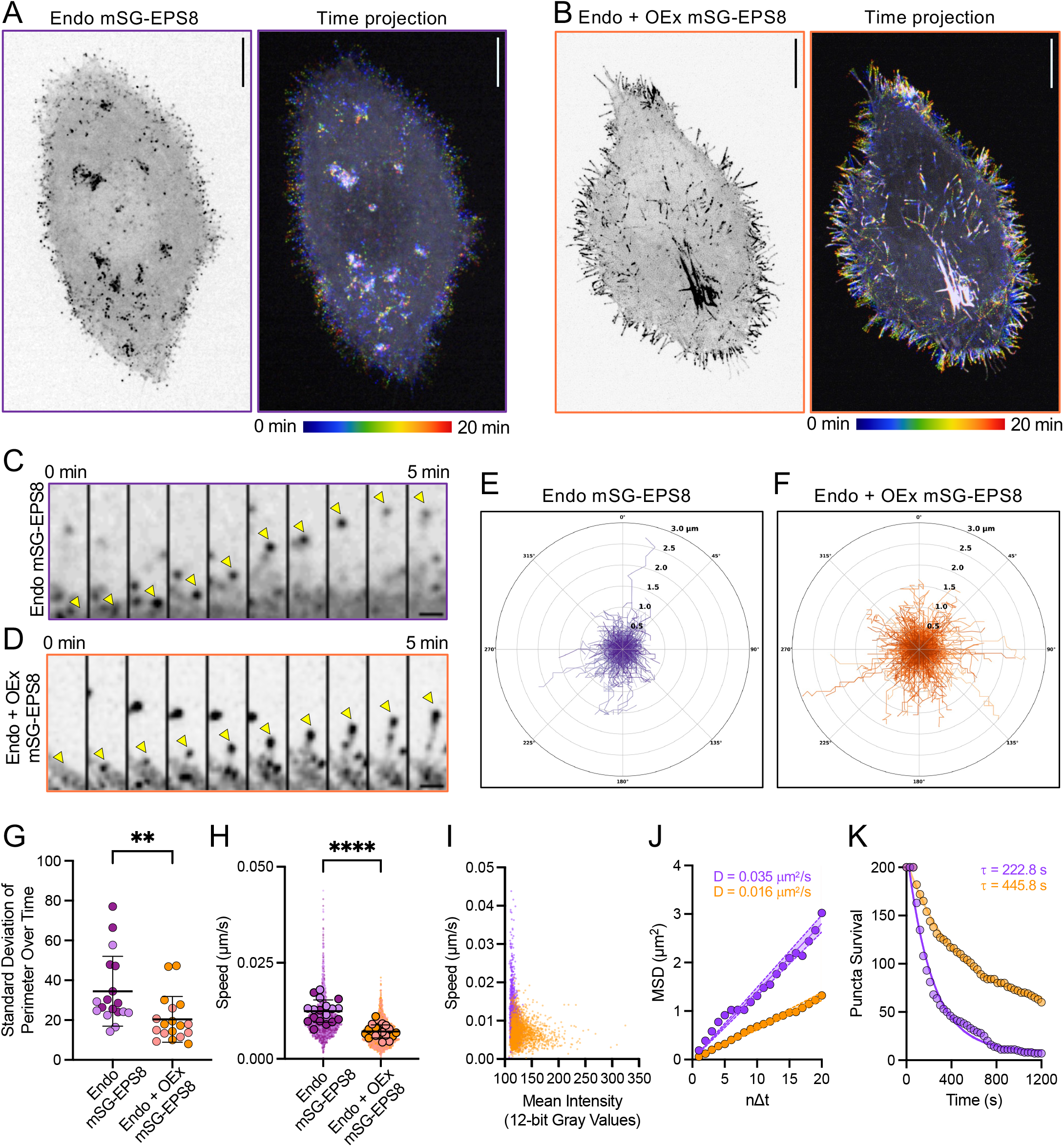
EPS8 dampens the growth dynamics and increases the lifetime of filopodia. **A - B)** (Left) MIP 100x SDC images of Endo-mSG-EPS8 (purple outline) and Endo + OEx mSG-EPS8 (orange outline) cells. Scale bars = 10 µm. (Right) Temporally color-coded projections of 11 frames from cells imaged over 20 min. **C - D)** MIP montage showing tip puncta in SDC timelapse of Endo-mSG-EPS8 (purple outline) and Endo + OEx mSG-EPS8 (orange outline) cells over 5 min. Arrows indicate filopodial growth events. Scale bar = 1 µm. **E - F)** Representative radial trajectory plots showing all puncta from 20 min of observation from cells shown in A (756 puncta) and B (1,315 puncta), respectively. Concentric circles start at 0.5 µm and extend to 3 µm in 0.5 µm increments. **G)** Standard deviation of whole cell perimeter changes over 20 min, from Endo-mSG-EPS8 vs. Endo + OEx mSG-EPS8 cells. 18 cells quantified from 2 biological replicates. Unpaired Mann-Whitney t-test, **p = 0.007. **H)** Superplots showing quantification of EPS8 puncta speeds measured from Endo-mSG-EPS8 (purple) vs. Endo + OEx mSG-EPS8 (orange) cells. Small, transparent data points show individual puncta and larger, opaque points show per cell averages. Unpaired t-test on per cell averages, ****p < 0.0001. **I)** Correlation plot of individual punctum mean 12-bit gray intensity values vs. speeds from Endo-mSG-EPS8 (purple points) and Endo + OEx mSG-EPS8 (orange points) cells. For H and I, 1,789 Endo-mSG-EPS8 puncta and 2,142 Endo + OEx mSG-EPS8 puncta were quantified over 18 and 19 cells, respectively, from 2 biological replicates. **J)** Mean-squared displacement (MSD) analysis of 200 puncta tracks from 18 Endo-mSG-EPS8 (purple) vs. 19 Endo + OEx mSG-EPS8 (orange) cells, from 2 biological replicates. **K)** Puncta survival analysis of 200 puncta from 18 Endo-mSG-EPS8 (purple) and 19 Endo + OEx mSG-EPS8 (orange) cells imaged for 20 min, every 30 sec, from 2 biological replicates.

### EPS8 dampens the rate of actin core bundle turnover

Filopodial dynamics are driven in large part by the assembly and disassembly of actin filaments that comprise the core bundle. Coordination of these activities leads to “treadmilling”, where new actin subunits enter filament barbed ends at the distal tips, flux retrograde through the bundle, and leave the bundle from the basal end (44). With this in mind, we asked if EPS8 supplementation dampens filopodia dynamics by slowing F-actin turnover in core actin bundles. To test this, we transiently expressed mScarlet3-β-actin in Endo mSG-EPS8 cells, with and without supplemental EPS8 (Fig. 4). Using total internal fluorescence (TIRF) microscopy, we were able to visualize fiducial patterns of F-actin retrograde flow over time. Using lines drawn along the filopodial axis, we generated kymographs and calculated retrograde flow velocities from the slopes of fiducial patterns in these images (Fig. 4A,B). Notably, in cells expressing supplemental EPS8, the retrograde flow rates were ∼5-fold lower relative to control cells expressing endogenous levels (Fig. 4C). This result suggests that EPS8 puncta dampen filopodial dynamics by slowing core bundle actin turnover.

**Figure 4.**
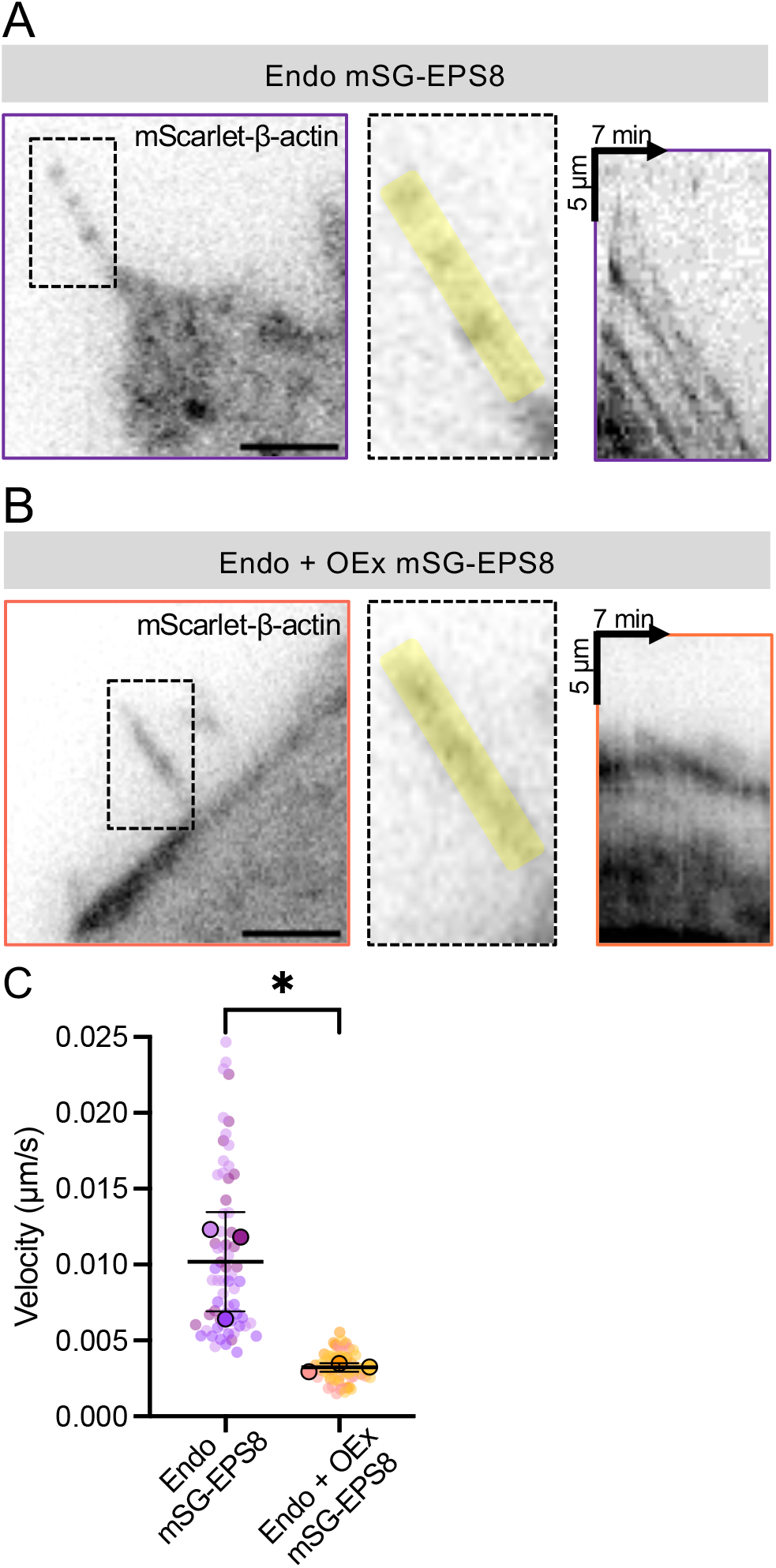
EPS8 reduces actin retrograde flow rates. **A - B)** Representative filopodia from 100X SDC TIRF images of (A) Endo-mSG-EPS8 cells and (B) Endo + OEx mSG-EPS8 cells overexpressing mScarlet3-β-actin. Cells were imaged for 7 min every 10 sec. Scale bar = 5 µm. Kymographs were drawn along the length of the highlighted yellow region. Box height = 5 µm, box width = 7 min. **C)** Superplots show quantification of the velocity of actin treadmilling as determined from kymographs described in A and B from Endo-mSG-EPS8 (purple) vs. Endo + OEx mSG-EPS8 (orange). Small, transparent data points represent the average of all kymographs taken per individual filopodium and larger, opaque points show replicate averages. 1-4 velocity measurements per kymograph were averaged from 2-4 filopodia per cell, quantified over 27 cells from 3 biological replicates. Unpaired t-test on the 3 biological replicates, *p = 0.0212.

### EPS8 protects core bundle filament barbed ends from Cytochalasin D

Cytochalasin D (Cyto D) is a toxin that strongly binds to the barbed ends of growing actin filaments, prevents the addition of new monomers, and leads to filament depolymerization (45). Previous work from our group showed that CytoD treatment reduced EPS8 distal tip localization in epithelial microvilli, raising the possibility that EPS8 binds directly to actin filament barbed ends (31). To further test this idea, we sought to determine if supplemental EPS8 could protect filopodial core bundles from Cyto D capping and disassembly. Here, we used SDC microscopy to image Endo mSG-EPS8 cells with or without supplemental EPS8 cells, before and after treatment with 500 nM Cyto D (Fig. 5A,B). As expected, filopodia in Endo mSG-EPS8 cells rapidly collapsed following the Cyto D treatment; they also experienced a coincident loss of distal tip EPS8 puncta (Fig. 5A). Counting the number of EPS8 puncta at the tips of filopodia pre- and post-Cyto D treatment revealed that the percentage of puncta remaining after treatment was significantly higher in cells with supplemental EPS8 vs. controls (Fig. 5C). These findings suggest that higher levels of EPS8 compete with Cyto D and can protect barbed ends against filament capping and disassembly.

**Figure 5.**
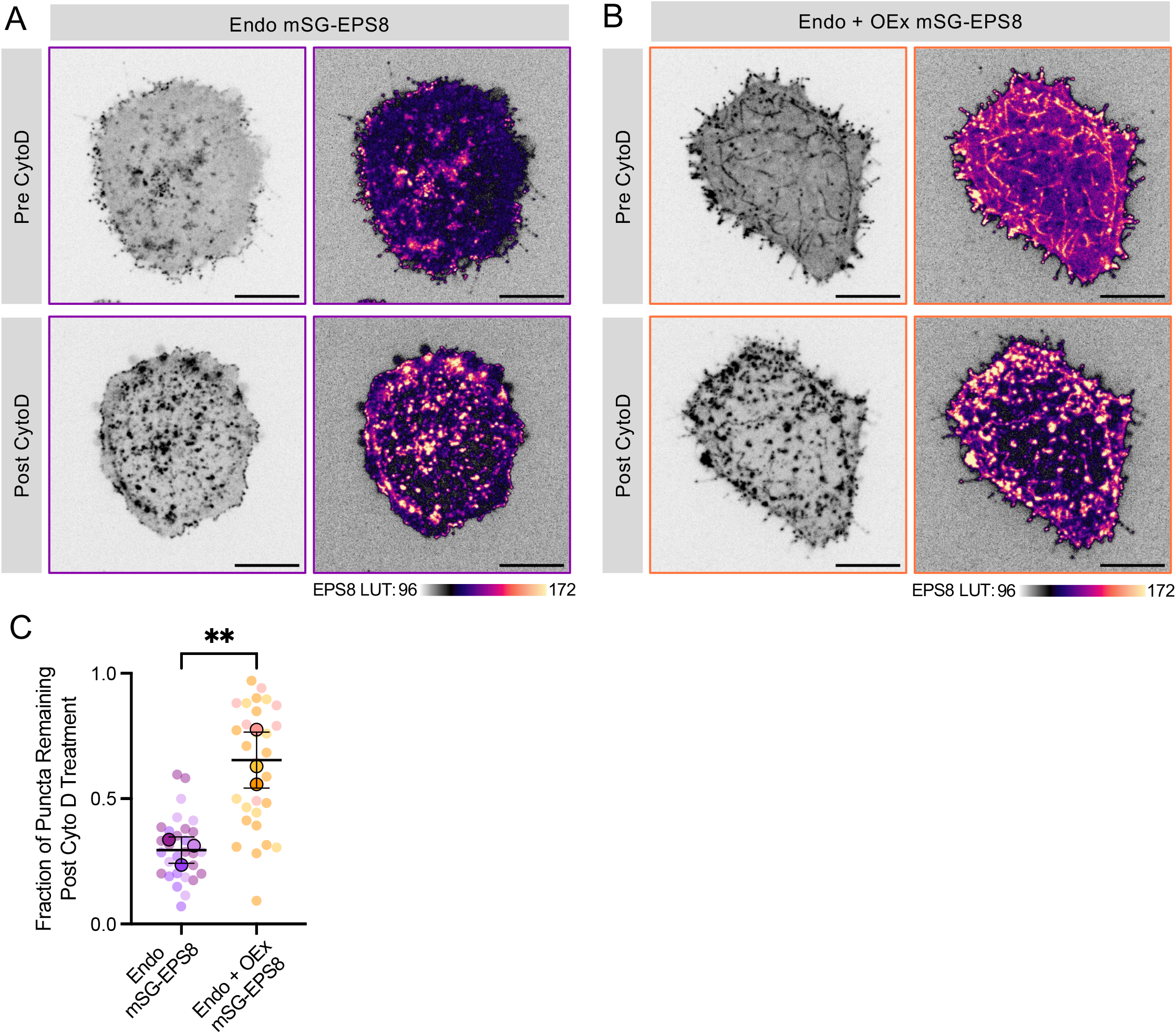
EPS8 protects filopodia from Cytochalasin D induced disassembly. **A-B)** MIP 100x SDC images of (A) Endo-mSG-EPS8 HeLa cells and (B) Endo + OEx mSG-EPS8 cells before and 10 min after 500 nM Cytochalasin D treatment. Scale bar = 10 µm. LUTs are intensity-matched and shown below images for A and B. **C)** Superplots show quantification of the percentage of EPS8 puncta remaining at the tips of filopodia starting 5 min after 500 nM Cytochalasin D treatment. Small, transparent data points show individual cells and larger, opaque points show replicate averages. Puncta were analyzed from 30 cells from 3 biological replicates. Unpaired t-test on the 3 biological replicates, **p = 0.0073.

### EPS8 and actin levels increase during intestinal brush border maturation

Previous studies revealed that EPS8 localizes to the tips of mature microvilli on the surface of fully differentiated enterocytes (15). Other studies also found EPS8 signal in the crypt compartment, where microvilli actively grow on the surface of differentiating enterocytes (31). If EPS8 promotes the growth and stability of protrusions, one might expect its levels to increase from the crypt to the villus, as more protrusions are assembled to build a functional brush border. To test this, we immunostained sections of mouse small intestine to directly compare EPS8 localization and intensity in differentiating vs. mature epithelial cells, in the crypt and villus compartments, respectively (Fig. S1A,B). For these experiments, we used paraffin embedded sections, which offered more robust staining relative to other tissue preparations (e.g. frozen sections in OCT). We also co-stained samples with a fluorophore conjugated γ-actin antibody, to label F-actin. As expected, image volumes revealed that γ-actin signal was significantly higher in villus vs. crypt epithelial cells, which is consistent with increased numbers of microvilli found on the villus. When EPS8 intensity was normalized to the γ-actin signal per enterocyte, we observed a ∼1.8-fold increase from the crypt to the villus (Fig. S1D); i.e. per unit F-actin, mature microvilli on the surface of villus enterocytes exhibit higher levels of EPS8 at their distal tips. These data further support a role for EPS8 in promoting the growth and stability of actin bundle-supported protrusions.

## DISCUSSION

Based on extensive cell biological studies, we now recognize a growing collection of proteins that target specifically to the distal tips of protrusions and are well positioned to control the elongation and stability of these structures. In filopodia, enabled/vasodilator-stimulated phosphoprotein (Ena/VASP) family proteins localize to the tips, where they serve to antagonize the activity of capping protein (CP), which would otherwise inhibit filament elongation (46). Other studies suggest roles for Ena/VASP in clustering filament barbed ends at these sites (47). Formins are tip-targeting actin filament nucleators that might also exhibit anti-capping activity and promote filopodial elongation (48). IRSp53 is a tip-localized factor studied primarily in filopodia, which may link growing actin filaments to highly curved regions of plasma membrane using its N-terminal I-BAR domain (49). Other I-BAR proteins, including IRTKS and Pinkbar, have also been localized to the tips of epithelial microvilli (17) and stereocilia (50), where they may play similar roles. In contrast to the apparent diversity of protein machinery noted in these different protrusion contexts, EPS8 is a rare common feature, exhibiting robust distal tip localization in all the cases studied to date (15,16,18). Despite this point, the specific activities that EPS8 brings to the tip compartment have remained unclear for many years. In this paper, we sought to clarify our understanding of how EPS8 functions in protrusion growth and maintenance. Our findings revealed that EPS8 directly increases the number of filopodia (Fig. 1 and Fig. 2H) and the F-actin content per protrusion (Fig. 2E-G), while simultaneously attenuating filopodial dynamics, prolonging protrusion lifetime (Fig. 3 and Fig. 4), and preventing the capping of barbed ends (Fig. 5).

Genetic perturbation experiments in multiple model systems revealed that common features of EPS8 loss-of-function (LOF) are shortened and malformed protrusions (15,27,29). However, protrusions do still form in these systems, and the severity of observed phenotypes might be limited by compensation with other members of the EPS8-like protein family: EPS8L1, EPS8L2, and EPS8L3 (51). EPS8L1 and EPS8L2 have the highest homology to EPS8, have been shown to localize to fibroblast ruffles, and are sufficient enough to restore ruffle formation in *Eps8*-null cells (40,51). EPS8L2 has the highest overlap of expression with EPS8 at the mRNA level, whereas EPS8L1 and EPS8L3 are more restricted to certain tissues such as the colon and small intestine (40). To avoid compensation due to upregulation of EPS8-like proteins that may occur in long-term LOF models, we leveraged an acute sequestration strategy to rapidly deplete EPS8 from the distal tips of filopodia. Based on the striking loss of EPS8 tip puncta and parallel loss of filopodia (Fig. 1), our findings clearly demonstrate that EPS8 is required for filopodia survival on the cell surface. Why are filopodia lost following EPS8 depletion? One possibility is that, in the absence of EPS8, filament barbed ends at the distal tips become capped and core bundles shorten, driven by turnover mechanisms that are still active at the basal ends. Based on the results of our live imaging and drug perturbation experiments, we consider this proposal in more detail below.

EPS8 was assigned capping and bundling functions based on early biochemical characterization of purified domains derived from its C-terminus (a.a. 648–821) (26,33). Capping activity was localized to an amphipathic helix containing key residues V689 and L693, whereas a more C-terminal four-helix bundle was found to exhibit filament bundling activity (33). Consistent with a role in bundling, we found that providing cells with supplemental EPS8 led to a parallel increase in the number of filaments per filopodial core bundle (Fig. 2). How EPS8 exerts its bundling potential from the distal tip compartment remains unclear, but there is precedent for this type of activity from studies of other tip targeting molecules such as VASP (52). Moreover, previous studies suggest that binding partners might tune this activity. Indeed, EPS8 function is activated by binding to Abi1, an interaction that relieves auto-inhibition and promotes actin-cable formation (26,53). Furthermore, when in complex with the membrane-binding I-BAR domain protein IRSp53, EPS8 displays strong filament cross-linking activity (27,30).

While IRSp53 is not enriched in stereocilia or epithelial microvilli, these structures do contain other similar I-BAR proteins. For example, epithelial microvilli contain IRTKS, which localizes to the distal tip compartment and uses its SH3 domain to bind directly to polyproline motifs in EPS8 (17). Whether the interaction between EPS8 and IRTKS stimulates bundling activity and/or controls filament number in microvillar core bundles has yet to be tested.

Beyond a role in bundling, our results suggest that EPS8 also functions directly at filament barbed ends to control G-actin incorporation. Indeed, increasing levels of EPS8 slowed the dynamics of filopodia growth, prolonged protrusion lifetimes, and reduced actin treadmilling speeds in these structures (Figs. 3 and 4). Additionally, higher levels of EPS8 protected core bundles against Cyto D (Fig. 5). Because this compound binds specifically to filament barbed ends, where it would typically inhibit elongation and induce disassembly, our results suggest the EPS8 is also a barbed end binder (Fig. 5). Based on these findings, we propose that EPS8 functions as a “leaky capper” that dwells at core bundle barbed ends, allowing the incorporation of new actin monomers at a submaximal rate, while preventing strong filament capping by *bona fide* capping protein (35), potentially through steric inhibition. This model would be similar to that proposed for formin family members, such as yeast Bni1p, which slows G-actin incorporation while protecting filaments from capping (54). Such a mechanism would also explain the outcome of our acute depletion experiments (Fig. 1), where the striking loss of filopodia following rapid sequestration of EPS8 is consistent with strong barbed end capping following this perturbation.

In summary, we leveraged acute loss-of-function and titrated gain-of-function approaches in combination with live imaging, to probe the function of EPS8 at the distal tips of filopodia. Our results provide clear evidence that this factor exhibits bundling potential and functions directly at core bundle filament barbed ends, to dampen protrusion growth dynamics, protect filaments from capping, and importantly, prolong protrusion lifetime. Based on these activities, EPS8 appears to function as a protrusion “survival factor”, which is necessary and sufficient for driving the accumulation of large numbers of protrusions on the cell surface. This role is likely critical for the formation of apical specializations, which contain 100s-1000s of long-lived protrusions that are essential for the physiological function of solute transporting and mechanosensory epithelia.

## EXPERIMENTAL PROCEDURES

### Lead contact

Further information and requests for resources and reagents should be directed to and will be fulfilled by the lead contact.

### Materials availability

Plasmids generated in this study will be made available from the lead contact on request.

### Experimental model and subject details

#### Cell culture

HeLa cells were cultured at 37° C and 5% CO_2_ in Dulbecco’s Modified Eagle’s Medium (DMEM) (Corning #10-013-CV) with high glucose and 2 mM L-Glutamine supplemented with 10% fetal bovine serum (FBS) and L-Glutamine (Corning # 25-005-Cl).

#### Cloning and constructs

All PCR reactions were performed with Q5 High-Fidelity DNA Polymerase (NEB #M0491S) and amplified fragments were assembled via Gibson assembly (NEB #E2621S). pmScarlet3-β-actin was generated by tagging β-actin with the mScarlet3 sequence from pmScarlet3-C1 (Addgene #189753) using a NheI and XhoI restriction digest. pmStayGold-EPS8 was generated by inserting the mStayGold sequence into pEGFP-C1-EPS8 (17,32) using primers: 5’-

GCGCTACCGGTCGCCACCATGGTGTCTACAGGCGAGGAGC-3’ and 5’-ATTCGAGATCTGAGTCCGGACAGGTGGGCCTCCAG-3’ to amplify the mStayGold sequence; 5’-AGACCCTGGAGGCCCACCTGTCCGGACTCAGATCTCGAATCAC-3’ and 5’-GCTGGGTCGAATTCGCCCTTTTAGTGACTGCT-3’ to amplify the EPS8 sequence; and 5’-ATGAAGGAAGCAGTCACTAAAAGGGCGAATTCGAC-3’ and 5’-AGCTCCTCGCCTGTAGACACCATGGTGGCGACCGGTAG-3’ to amplify the plasmid backbone with mStayGold and EPS8 homology. pcDNA3.1Zeo-mStayGold-EPS8-FRB-Myc was generated by inserting mStayGold from pmStayGold-EPS8 into FRB-myc (Addgene #20228) using a HindIII-HF and XhoI restriction digest. Primers 5’-TGACTGAAGCTTGCCACCATGGTGTCTACA-3’ and 5’-CAGTCACTCGAGGCTTCCTCCGTGACTGCTTCCTTCATCAAA-3’ were used to amplify the mStayGold sequence with plasmid backbone homology. pmCherry-N3-TOMM20-FKBP was synthesized and purchased from VectorBuilder. All plasmids were confirmed by sequencing.

#### Endogenous tagging of EPS8 with mStayGold in HeLa cells

A guide RNA (gRNA) for targeting human EPS8 was generated using the Benchling CRISPR design tool. A single 20 bp gRNA with an NGG PAM site was selected based on the on-target score and guide RNA overlap with output from the GenScript gRNA program. gRNA sequences were adapted to include overhangs complementary to the PX459v2.0 backbone. Oligos were synthesized by Sigma-Aldrich and resuspended at a concentration of 100 µM. Following the Zhang laboratory Addgene protocols (https://www.addgene.org/crispr/zhang/), the PX459v2.0 plasmid was cut using BbsI and gel purified (Macherey-Nagel #740609.50). Forward and reverse oligos were phosphorylated, annealed, ligated into the cut PX459v2.0 vector, and subsequently transformed into DH5α bacteria. Plasmid DNA from positive clones was purified and sequenced using whole plasmid sequencing. A model repair construct encoding mStaygold plus a linker sequence, flanked by homology arms that target immediately upstream of the *Hs* EPS8 exon 2, was created *in silico* using the GenScript CRISPR design program. This sequence was used to design 100 bp forward and reverse primers, which contained 80 bp of genomic sequence, and 17 bp or 10 bp of mStayGold sequence, respectively (Forward primer, 5’GAAACTGAGATGTCTTGACCTTTTTATTGCCTTTAATCTATGTGTCTTTCTTTTTTTAGAACACACAAGTGAAAGACACAATGgtgtctacaggcgagga 3’, 80bp EPS8 homology arm, ATG added for start codon, and 17 bp mStayGold homology; Reverse primer, 5’AAACTTTTCAACTTACTTCATCTGAGATGGGTACATTCCAAAACTACTGGGATGATTAGAAATATGACCATTCATAGCGGAGGAGGAGGAcaggtgggcc 3’, 75 bp EPS8 homology arm, 15 bp SGGGG linker, and 10 bp mStayGold homology). A start codon was added to the forward primer and a linker sequence encoding SGGG to the reverse primer. PCR was performed using the Guide it Long ssDNA Strandase Kit V2 (Takara #532667) to generate double stranded DNA encoding the full repair construct containing the mStayGold sequence. This PCR product was confirmed by gel electrophoresis and subsequently purified (Macherey-Nagel #740609.50). Transfection of gRNA plasmid and double stranded repair construct DNA into HeLa cells was performed with Lipofectamine 2000 (Thermo Fischer #11668019) in a 6 well format. Cells were left in transfection reactions for two days, expanded into T175 flasks, and then sorted as single cells into 96 well plates by the Vanderbilt Flow Cytometry Core. Positive cells were identified by plating clones onto a glass bottom 6 well plate and imaging with a spinning disk confocal.

#### Transient transfections

Transfections were performed using Lipofectamine 2000 (Thermo Scientific #11668019) according to the manufacturer’s protocol. Cells were incubated in Lipofectamine and transfected with 100 ng - 2.2 µg total of DNA, overnight in 35mm glass-bottom dishes (Cellvis #D35–20–1.5-N), 4-chamber 35 mm glass-bottom dishes (Cellvis #D35C4-20-1.5-N), or on glass coverslips. Cells were then fixed or imaged live (more below).

#### F-actin labeling in live cells

SPY650-FastAct_X (Cytoskeleton #CY-CS502) was added to cell culture media at 1:1000 (Fig. 2) or 1:2000 (Fig. 1); cells were then incubated for 30 min - 1 hr before labeling media was removed and replaced with fresh media.

#### Immunofluorescence staining

*For cell staining:* Cells were rinsed in prewarmed PBS and fixed by incubating in prewarmed 4% paraformaldehyde (Electron Microscopy Services #15710) for 15 min at 37° C. Cells were then rinsed 3x for 5 min with PBS and permeabilized in 0.1% Triton X-100 (Sigma #T8787) for 15 min at RT. Cells were then rinsed with PBS 3x for 5 min and then blocked in 5% BSA for 1 hr at 37°C. Following blocking, cells were briefly rinsed 1x with PBS and then incubated with 1° antibody (rabbit polyclonal anti-non muscle myosin IIA; BioLegend #909802) at 1:200 for 1 hr at 37° C. Cells were then rinsed 4x for 5 min with PBS and incubated with 2° antibodies (Goat-anti-Rabbit Alexa Fluor 647; Invitrogen, #A21246) at 1:1000 and Alexa Fluor Plus 405 Phalloidin (Thermo Scientific #A30104) at 1:200 dilution for 1 hr at RT. Following 2° antibody incubation, cells were rinsed 4x for 5 min in PBS and mounted on glass slides using ProLong Gold antifade reagent (Life Technologies). *For tissue staining:* Paraffin-embedded tissue sections of mouse small intestine were deparaffinized using HistoClear II solution (National Diagnostics #HS-202) twice for 3 min. Tissue was then rehydrated through a descending ethanol series (100%, 95%, 90%, 70%, 50%) and subject to antigen retrieval in a boiling buffer (10 mM Tris, 0.5 mM EGTA) at pH 9 for 1 hour. After cooling, sections were washed and blocked with 10% normal goat serum (NGS) for 1 hr at RT and stained with primary antibody (mouse anti-EPS8; BD Transduction Laboratories, #610144, 1:400; or γ-actin 1-17 Alexa Fluor 647, Santa Cruz #sc-65638, 1:100) overnight in 1% NGS at 4° C. The next day, slides were washed 3x, 5 min each in PBS, then incubated with secondary antibodies at 1:1000 (Goat-anti-mouse Alexa Fluor 488; Molecular Probes, #A11031), diluted in 1% NGS for 1 hr at RT in the dark. Slides were then washed 3x, 5 min each in PBS followed by dehydration with an ascending ethanol series (50%, 70%, 90%, 95%, 100%, 100%) 5 min each. Coverslips were mounted with ProLong Gold antifade reagent (Life Technologies).

#### Drug Treatments

To induce heterodimerization of FRB- and FKBP-tagged fusion proteins expressed in HeLa cells (Fig. 1), cells were imaged prior to the addition of 500 nM of A/C Heterodimerizer (Takara #635057), and again after 1 hr. For the drug treatments (Fig. 5), cells were imaged prior to the addition of 500 nM of cytochalasin D (Sigma #C2618), and again after 5-10 min.

#### Light microscopy, acquisition, and image processing

##### Microscopy

Live imaging for Figs. 2, 3, and 5 was performed on a Nikon Ti2 inverted light microscope equipped with a Yokogawa CSU-X1 spinning disk, 488 nm and 647 nm excitation LASERs, an Apo TIRF 100x/1.49 NA oil objective, and a Photometrics Prime 95B sCMOS camera. For Figs. 1 and 4, and S2, live imaging or fixed imaging was performed on a Nikon Ti2 inverted light microscope equipped with a Yokogawa CSU-W1 spinning disk head, with 488 nm, 561 nm, and 647 nm excitation LASERs, an Apo TIRF 100x/1.49 NA oil objective, and a Photometrics Prime 95B sCMOS camera SDC (Fig. 1 and S2) or TIRF optics on a separate port using a Hamamatsu Fusion BT (Fig. 4). Live cells were maintained in a stage top incubator at 37 °C with 5% CO2 (Tokai Hit). For Fig. S1, fixed imaging was performed using a Nikon AX/AXR confocal microscope, 488 nm and 647 nm excitation LASERs, a Plan Apo λD 100x/1.45 NA Oil objective, and an NSPARC detector. *Acquisition:* For all timelapse imaging, 0.5% methylcellulose was added to media prior to imaging to reduce lateral waving of filopodia. Z-stacks of ∼1-4 µm with a step size of 0.2 µm was acquired for live and fixed cells, depending on the intended analysis, using a triggered NIDAQ Piezo stage. Tissue samples were imaged at 1024 x 1024 resolution, and 73 z-stacks were collected with a step size of 0.1 μm.

##### Processing

All 3D images and movies were deconvolved using Nikon Elements. For figure preparation, look up tables were optimized to facilitate visualization. All images and movies in the same experiment, compared across conditions and in each replicate, were collected using matched laser powers and camera exposure times.

### Quantification and statistical analysis

All images were processed and analyzed using Nikon Elements software or FIJI. Analysis was performed on raw images, except for the crypt and villus intensities and filopodia counting analyses where images were first deconvolved. Analysis was done on maximum intensity projection (MIP) or sum intensity projection (SIP), except for the retrograde flow analysis.

#### Analysis of filopodia number

Filopodia were manually annotated using Nikon Elements “Count” function, using the F-actin signal channel.

#### Analysis of EPS8 and γ-actin intensity and the dynamics of EPS8 puncta

For the analysis of EPS8 and γ-actin signal on cells of the crypt and villus, drawing tools in FIJI were used to create an ROI along the apical surface of a cell in the γ-actin channel using a sum-intensity projection. The ROI was then saved and applied to the same cell in the EPS8 channel to obtain the apical EPS8 intensities. For analysis of whole cell intensities, a binary was drawn around each cell in Nikon Elements, and a threshold was set on the signal. For measurements of EPS8 puncta intensities and dynamics, a toroidal binary was drawn around the peripheral filopodia excluding the cell bodies and Nikon Elements “Bright Spot Detection” was used to threshold the signal of puncta within the binary. Dynamic data were collected with the “Track Binaries” function. Dynamic measurements excluded puncta lasting less than 3 frames or puncta that were not present at the beginning of the movie.

#### Area under the curve (AUC) analysis

In FIJI, we generated line scans (5 pixels in width, 1.4 µm in length) orthogonal to individual protrusions visualized in the FastActX channel. Line scans were drawn to ensure lines only crossed through one protrusion per line (42). Since filopodia are not uniform in width, line scans were drawn across the midpoint to standardize measurements. Areas under the resulting intensity curves were then calculated in PRISM v.10, averaged, and plotted. The resulting metric is expected to reflect the F-actin quantity in each core bundle.

#### Radial trajectory plots

Representative radial plots were generated using (x,y) positions of puncta and directional headings over time. These data were obtained from the “Track Binaries” function in Nikon Elements and plotted using ChatGPT.

#### Mean-squared displacement analysis

EPS8 puncta trajectories, as an indicator for filopodia motion, were analyzed using mean-squared displacement (MSD) analysis (43). MSD curves were fit with a model describing diffusive motion in the form MSD(nΔt) = 4DnΔt, where D is the diffusion coefficient and nΔt is total time window. Curve fitting and sum-of-squares F-tests were performed using PRISM v.10.

#### Puncta survival analysis

To assess the survival probability of EPS8 puncta, the number of puncta present at each time point was manually calculated from the “Track Binaries” data. Curves were fit to a one-phase decay model Y = Plateau+(Y_0_-Plateau)e^−KX^, where Plateau is the largest value of Y, Y_0_ is the initial Y value at X=0, and e^−KX^ is the amplitude of decay. Curve fitting and sum-of-squares F-tests were performed using PRISM v.10. For the survival of puncta following Cyto D treatment, Nikon Elements was used to draw toroidal binaries around the peripheral filopodia of the cells, excluding the cell body. “Bright Spots Detection” was used to identify puncta at filopodial tips and the number of puncta were determined. The percentages of puncta remaining were manually calculated and the puncta survival analysis was done on each cell before and after Cyto D treatment.

#### Retrograde flow analysis

Movies were collected at 10 second intervals over 7 min. Kymographs were generated for individual filopodia using Nikon NIS-Elements software on the mScarlet3-β-actin signal. Filopodia velocity rates were quantified by measuring the slope of fiducial actin trajectories along the filopodial axis using the velocity analysis function in NIS-Elements. Instances of negative slopes were counted as retrograde flow. Data from individual filopodia were pooled for each cell.

#### Statistical analysis

All statistical analyses were performed using PRISM v.10 (GraphPad) using a paired t-test for pairwise comparisons, an unpaired t-test, or a Mann-Whitney t-test. All error bars represent mean ± SD. Number of samples, measurements, and biological replicates are reported in each figure legend. Exact statistical tests and significance are reported in each figure legend.

## ACKNOWLEDGMENTS

The authors would like to thank all members of the M.J.T. laboratory for their feedback and guidance. This work was supported by NIH grants R01-DK095811 and R01-DK111949 (M.J.T.). We also acknowledge the Lacks family and are grateful for the use of HeLa cells, which heavily contributed to the discoveries in this work.

