## Supporting Information for "EPS8 dampens the growth dynamics and prolongs the lifetime of actin-based protrusions"

**SUPPLEMENTARY FIGURES – Mulligan et al.**

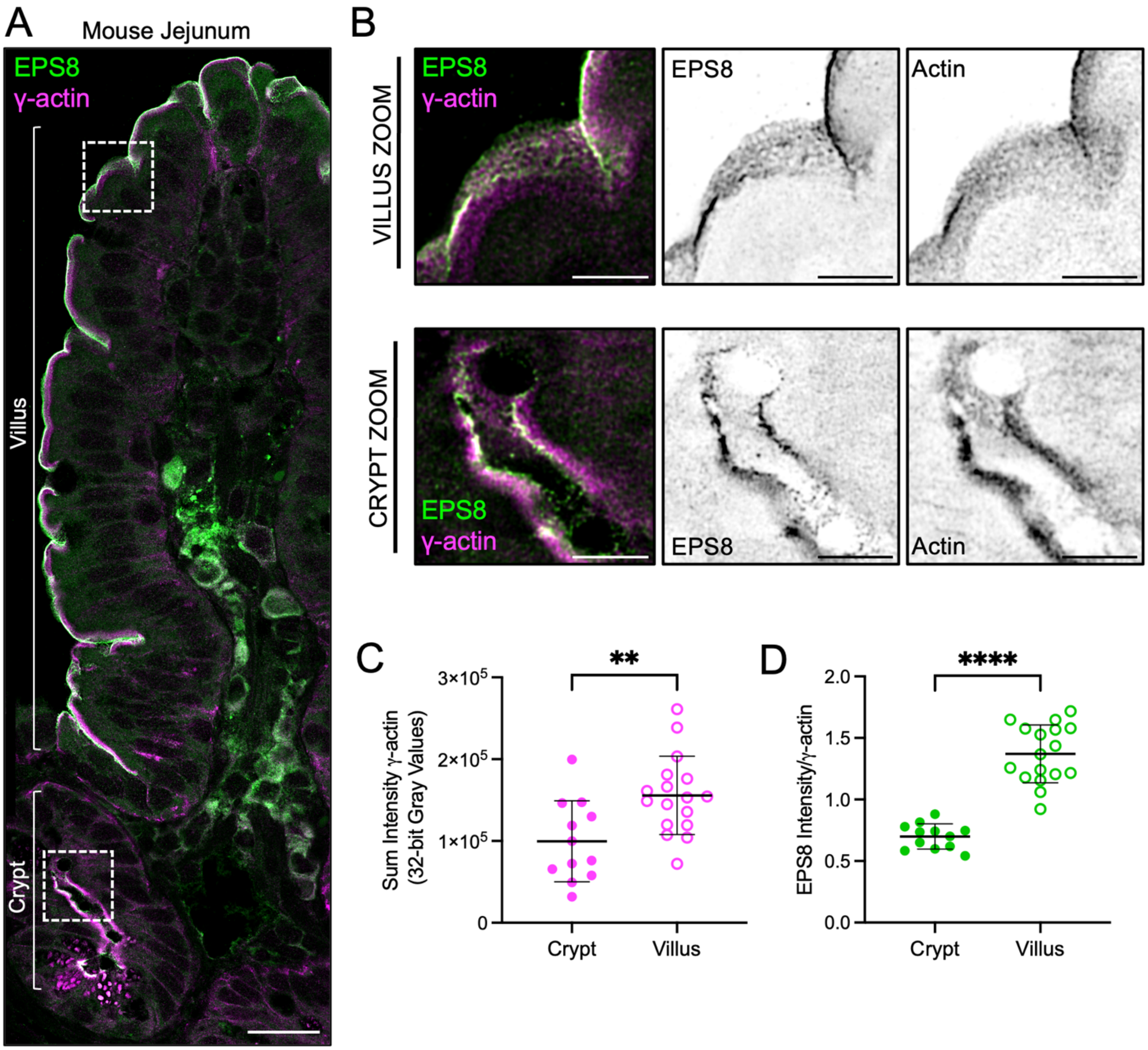


**Figure S1. EPS8 and actin levels increase during brush border maturation.**
**(A)** Stitched composite of 5 deconvolved 100X NSPARC images from the crypt to the villus in a small intestinal section stained for EPS8 (green) and γ-actin (magenta). Scale bar = 20 μm. **(B)** Zoom insets of the villus (top) and crypt (bottom) from (A). Each panel shows the merge with their inverted EPS8 and γ-actin channels. Scale bar = 5 μm. **(C)** Sum intensity 32-bit gray values of γ-actin in crypt vs villus cells. 12 and 17 cells were analyzed, respectively. Unpaired Mann-Whitney t-test, $p = 0.0043$. **(D)** Fluorescence intensity of EPS8 in crypt vs villus cells relative to the γ-actin intensity as shown in (C). 12 and 17 cells were analyzed, respectively. Unpaired Mann-Whitney t-test, $p = <0.0001$.

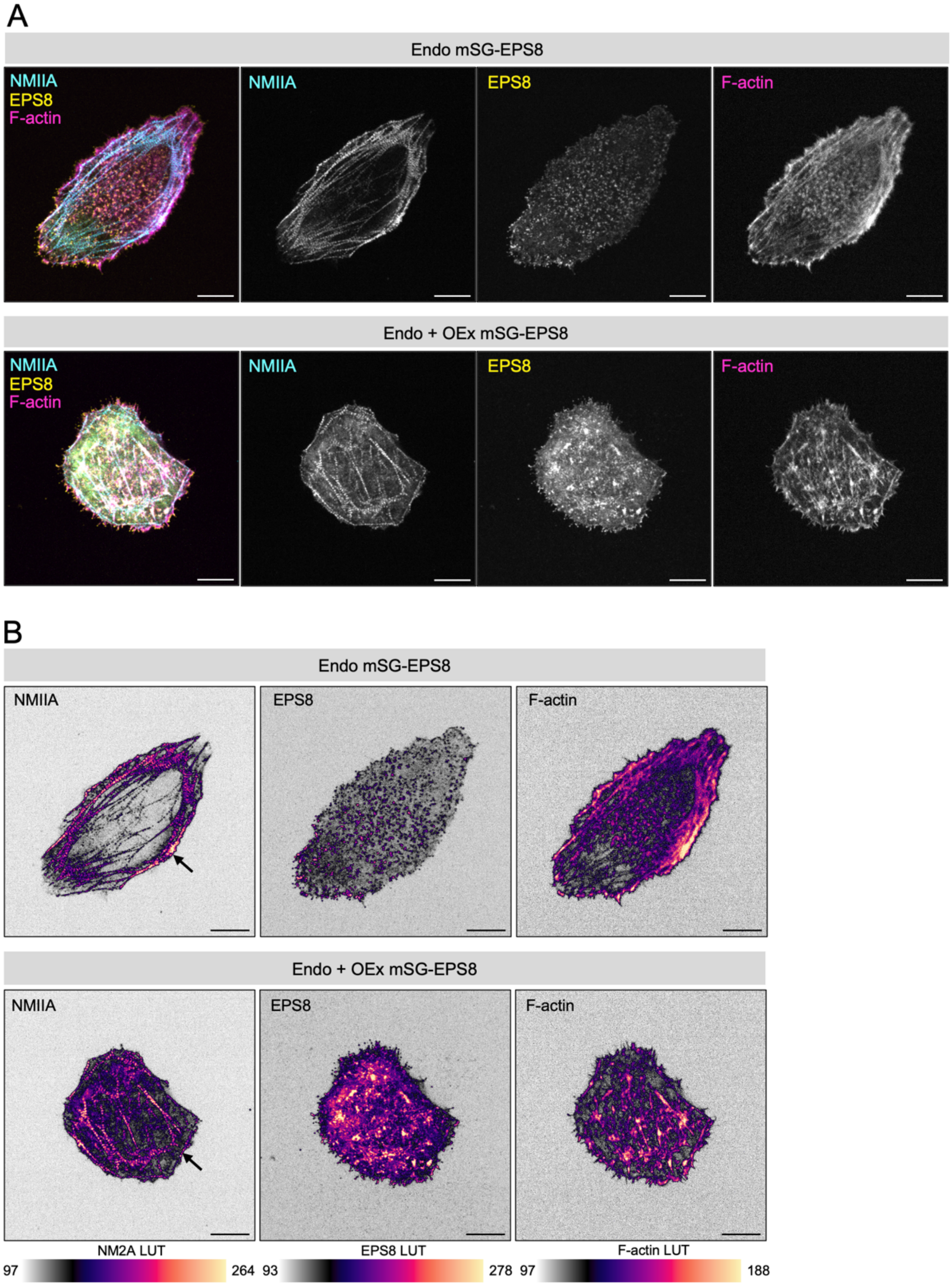


**Figure S2. Cortical NMIIA levels decrease when filopodial tip puncta EPS8 levels increase.**
**(A)** MIP 100X SDC images of Endo-mSG-EPS8 (top, yellow) Endo + OEx mSG-EPS8 (bottom, yellow) stained for NMIIA (cyan) and F-actin with phalloidin (magenta). Each panel shows the merge with each individual single channel. Scale bar = 10 µm. **(B)** Inverted single channels from (A) with intensity matched LUT and LUT scale per channel. Black arrows point to cortical NMIIA signal. Scale bar = 10 µm.

**SUPPLEMENTARY VIDEO LEGEND – Mulligan et al.**

**Movie S1. Supplemental EPS8 dampens filopodia dynamics, related to Figure 3.**
Live cell movies of maximum intensity projections of Endo mSG-EPS8 and Endo + OEx mSG-EPS8 HeLa cells imaged every 30 sec over 20 min. MIPs are composed of 13 x 0.2 µm confocal slices. Scale bar = 10 µm.
